# Super-resolution imaging uncovers the nanoscopic segregation of polarity proteins in epithelia

**DOI:** 10.1101/2020.08.12.248674

**Authors:** Pierre Mangeol, Dominique Massey-Harroche, Fabrice Richard, Pierre-François Lenne, André Le Bivic

## Abstract

Epithelial tissues acquire their integrity and function through the apico-basal polarization of their constituent cells. Proteins of the PAR and Crumbs complexes are pivotal to epithelial polarization, but the mechanistic understanding of polarization is challenging to reach, largely because numerous potential interactions between these proteins and others have been found, without clear hierarchy in importance. We identify the regionalized and segregated organization of members of the PAR and Crumbs complexes at epithelial apical junctions by imaging endogenous proteins using STED microscopy on Caco-2 cells, human and murine intestinal samples. Proteins organize in submicrometric clusters, with PAR3 overlapping with the tight junction (TJ) while PALS1-PATJ and aPKC-PAR6β form segregated clusters that are apical of the TJ and present in an alternated pattern related to actin organization. CRB3A is also apical of the TJ and weakly overlaps with other polarity proteins. This organization at the nanoscale level significantly simplifies our view on how polarity proteins could cooperate to drive and maintain cell polarity.

## Introduction

In epithelial tissues, cells coordinate their organization into a polarized sheet of cells. Each cell acquires an apico-basal organization and specialized lateral junctions, namely tight junctions (TJs, also known as zonula occludens), adherens junctions and desmosomes (Farquhar & Palade, 1963). This organization is key to the development, the maintenance, and the function of epithelial tissues. How this organization is orchestrated remains largely unknown.

Over the past two decades, a number of proteins have been discovered to be pivotal to epithelial polarization, such as PAR3, PAR6 and aPKC (PAR complex), Crumbs, PATJ and PALS1 (Crumbs complex) and Scribble, LGL and DLG (Scribble complex) in mammals (for review see (Assémat et al., 2008; Pickett et al., 2019; Rodriguez-Boulan & Macara, 2014). These proteins are remarkably well conserved over the animal kingdom (Belahbib et al., 2018; Le Bivic, 2013). Deletion or depletion of one of these proteins usually results in dramatic developmental defects (Alarcon, 2010; Charrier et al., 2015; Hakanen et al., 2019; Lalli, 2012; Park et al., 2011; Sabherwal & Papalopulu, 2012; Tait et al., 2020; Whiteman et al., 2014).

In the quest to understand the role of polarity proteins, numerous genetic and biochemical studies have been carried out. We and others have found that these proteins interact to form multiprotein complexes. Pioneering studies defined three core complexes based on the discovery of protein interactions or localization: the PAR complex consisting of PAR3, PAR6, and aPKC proteins (Joberty et al., 2000; Lin et al., 2000), the Crumbs complex consisting of CRUMBS, PALS1, and PATJ (Bhat et al., 1999; Makarova et al., 2003; Roh, Makarova, et al., 2002), and the Scribble complex consisting of Scrib, Lgl, and Dlg (Bilder et al., 2000). However, this view became more complex over the years as many interactions between proteins of different complexes can occur (Assémat et al., 2008; Hurd et al., 2003; Lemmers et al., 2004), and interactions of polarity proteins with cytoskeleton regulators and lateral junction proteins are common (Assémat et al., 2008; Chen & Macara, 2005; Itoh et al., 2001; Médina et al., 2002; Michel et al., 2005; Roh, Liu, et al., 2002; Takekuni et al., 2003; Tan et al., 2020). A current limitation in the understanding of polarization is that there is no clear hierarchy in the importance of these numerous interactions. Potential interactions revealed through biochemical assays do not necessarily reflect relevant interactions in cells, and do not specify when nor where in the cell these interactions could be relevant.

Polarity proteins have been localized with classical light microscopy and remarkably, they are often found concentrated at the apical junction, a key organizational landmark of epithelial cells. To understand how polarity proteins cooperate to orchestrate cell polarization, one needs to understand how precisely polarity proteins organize with respect to apical junctions or to the cytoskeleton. However, except from a few limited cases (Hirose et al., 2002; Izumi et al., 1998; Tan et al., 2020), the precise localization of polarity proteins at these organizational landmarks is missing. Moreover, knowing how polarity proteins organize in relation to each other in the cell should enable us to decipher from their plentiful known potential interactions, which ones are more relevant in specific sub-regions of the cell.

To tackle these challenges, we decided to systematically localize with STED microscopy, the polarity proteins that are key to the establishment of the apical pole of epithelia: PAR3, aPKC, PAR6β, PATJ, PALS1 and CRB3A. These proteins localize at the apical junction region of epithelial cells. Because how proteins interact and localize is likely to depend on cell differentiation, we decided to focus here on mature epithelia, a state where we hypothesize that protein interactions and localization are stationary. Using human and murine intestine and Caco-2 cells, we first imaged endogenous polarity proteins with respect to the TJ, to appreciate their overall organization in the region. Second, we localized these proteins two-by-two, to uncover relevant apical polarity protein sub-cellular associations. Finally, we focused on polarity proteins organization with respect to the actin cytoskeleton. We find that polarity proteins localize in distinct sub-regions that do not reflect the canonical definition of polarity proteins complexes. In addition, their localization with respect to the cytoskeleton emphasizes some emerging roles of polarity proteins as regulators of actin organization.

## Results

### Polarity proteins are localized in separate subdomains in the apical junction region

To obtain a first estimate of polarity protein localization in the TJ region, we systematically imaged each polarity protein with respect to a marker of the TJ. To this end, each apical polarity protein and a tight junction marker (ZO-1 or occludin) were immunostained and imaged together using Stimulated-Emission-Depletion (STED) microscopy (Hell & Wichmann, 1994) (Figure 1 and 2). STED images were acquired in the TJ region both in the apico-basal and the planar orientations of cells in human and mouse intestinal biopsies (Figure 1) and Caco-2 cells (Figure 2). To optimize the sample orientation, samples were cryo-sectioned when needed, in particular to obtain apico-basal orientation. Since we focused on mature epithelia, intestinal cells where observed exclusively in villi and Caco-2 cells were seeded on filters and grown over 14 days to allow differentiation (Pinto et al., 1983). Because the resolution of STED microscopy followed by deconvolution was, in our hands, about 80 nm in each color channel, the gain of resolution compared to classical confocal microscopy approaches was 3-fold in the planar orientation, and 7-fold along the apico-basal axis.

**Figure 1.**
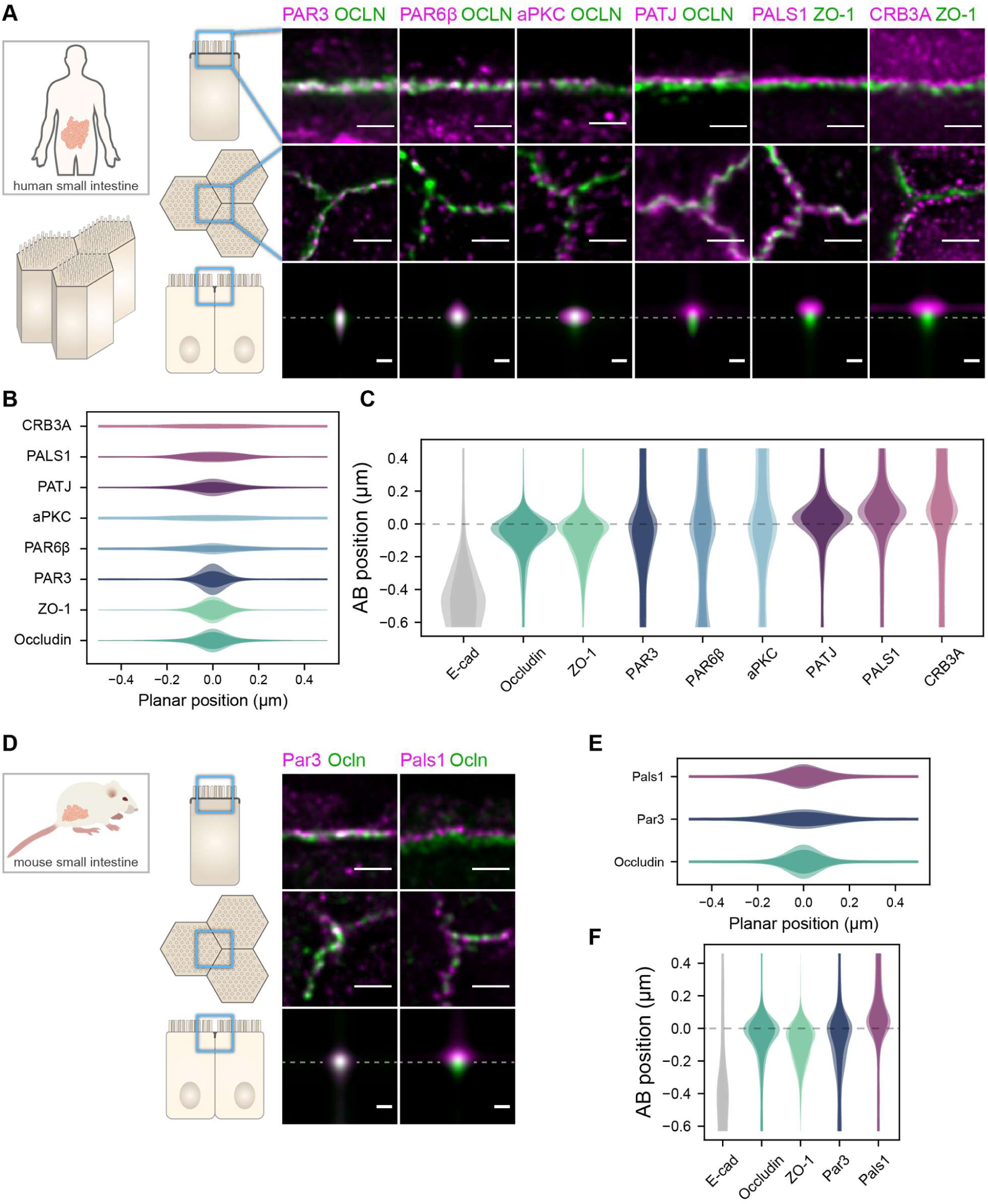
Polarity proteins localize in separate subdomains in the TJ region in human (A-C) and murine (D-F) small intestine biopsies. (**A**,**D**) STED images of protein localization in the TJ area. TJ proteins in green, polarity proteins in magenta. Top row, apico-basal orientation. Middle row, planar orientation. Bottom row, estimate of average protein localization in the apico-basal orientation perpendicular to the junction, obtained by multiplying average localizations estimated in (B) and (C) for human biopsies and (E) and (F) for murine biopsies. Top row and middle row, scale bar 1 µm; bottom row scale bar 200 nm. (**B**,**E**) Average localization of polarity proteins in the planar orientation, obtained by measuring the intensity profile of proteins perpendicular to the junction, using the TJ protein position as a reference. (**C**,**F**) Average localization of polarity proteins in the apico-basal orientation, obtained by measuring the intensity profile of proteins along the apico-basal orientation, using the TJ protein position as a reference. In (B,C,E,F), on a given position dark colors represent average intensity values, and lighter colors the average added with the standard deviation. The number of junctions used in quantification and details of the analysis are specified in the Material and Methods section.

**Figure 2.**
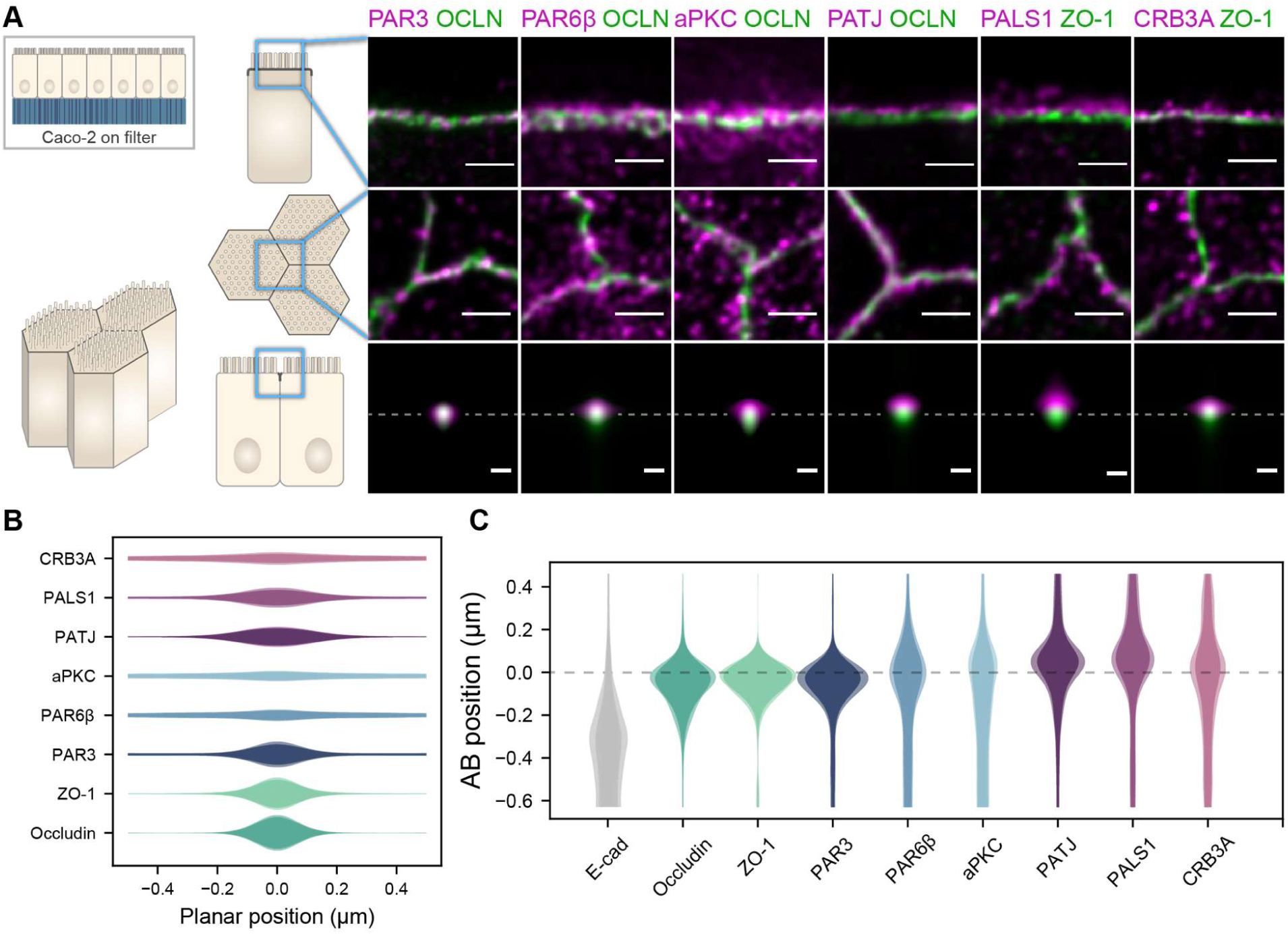
Polarity proteins localize in separate subdomains in the TJ region in Caco-2 cells. (**A**) STED images of protein localization in the TJ area. TJ proteins in green, polarity proteins in magenta. Top row, apico-basal orientation (obtained from cryo-sectioning cells grown on filter). Middle row, planar orientation. Bottom row, estimate of average protein localization in the apico-basal orientation perpendicular to the junction, obtained by multiplying average localizations estimated in (B) and (C). Top row and middle row, scale bar 1 µm; bottom row scale bar 200 nm. (**B**) Average localization of polarity proteins in the planar orientation obtained by measuring the intensity profile of proteins perpendicular to the junction, using the TJ protein position as a reference. (**C**) Average localization of polarity proteins in the apico-basal orientation obtained by measuring the intensity profile of proteins along the apico-basal orientation, using the TJ protein position as a reference. In (B,C), on a given position dark colors represent average intensity values, and lighter colors the average added with the standard deviation.The number of junctions used in quantification and the details of the analysis are specified in the Material and Methods section.

We found that the localization of each polarity protein was conserved across all samples and species (Figure 1 and 2). All proteins were concentrated in the TJ region as clusters of typically 80 to 200 nm in size (the smallest cluster size found is likely due to the imaging resolution limit), but their precise localization was protein dependent. We could group proteins in three main localization types. While we mostly found PAR3 at the TJs (Figure 1A,C,D,F and 2A,C), PAR6β and aPKC were at the TJ level and apical of the TJ (Figure 1A,C and 2A,C). We found CRB3A, PALS1 and PATJ almost exclusively apical of the TJ (Figure 1A,C,D,F and 2A,C). Interestingly, we often found PAR6β, aPKC, CRB3A, PALS1 and PATJ separated laterally from the TJ, since we frequently detected clusters of these proteins 100 to 200 nm away from the TJ (Figure 1A,B,D,E and 2A,E). There were some slight differences between intestinal samples and Caco-2 cells that may originate from sample preparation or from differences in cell organization due to tissue maturation. These first results show that polarity proteins organize in separate subdomains in the TJ region, namely PAR3 at the TJ and the other polarity proteins studied mostly apical of the TJ.

### Redefining relevant interactions between polarity proteins from colocalization analysis

The organization of proteins in separate subdomains led us to investigate how polarity proteins were organized within these subdomains, and more specifically how clusters of polarity proteins were localized with respect of each other. To tackle this question, we imaged polarity proteins two-by-two in Caco-2 cells and quantified the extent of their colocalization, using the protein-protein proximity index developed in (Wu et al., 2010), providing a quantitative estimate of protein proximity (Figure 3). Because of the organization of protein clusters, different proteins that localize at the same level on the apico-basal axis may appear as overlapping “more” when observed in the apico-basal orientation rather than when they are observed in the planar orientation; this is due to the fact that the axial-resolution (about 550 nm) is 7-fold lower than the planar resolution (about 80 nm). To circumvent this limitation, we minimized the apparent colocalization for each protein pair by orienting our sample either in the planar or apico-basal orientation, wherever apparent colocalization was lowest.

**Figure 3.**
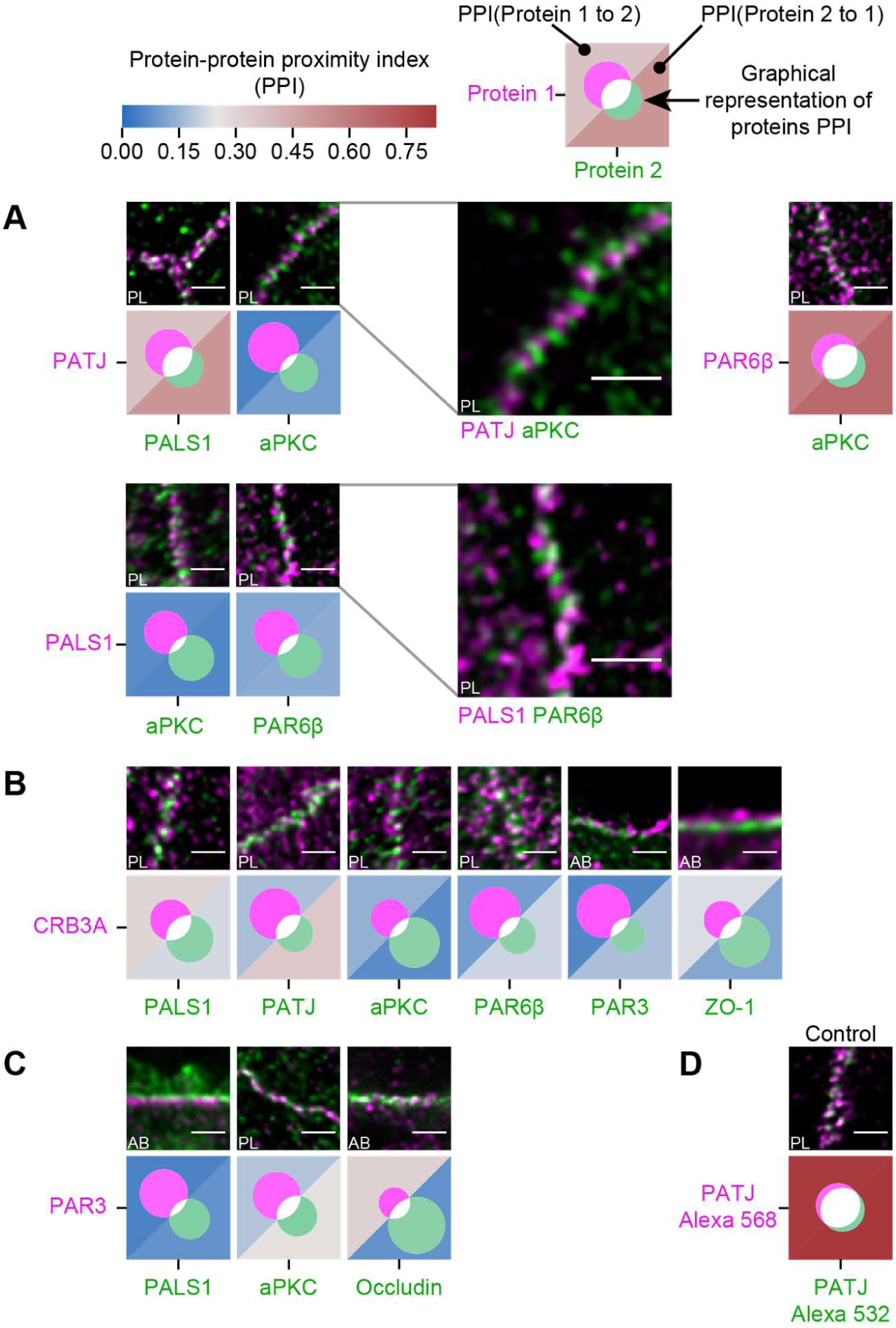
Proximity analysis of polarity proteins redefines protein complexes. The analysis is carried out in Caco-2 cells, where we used the concept of protein-protein proximity index (PPI) introduced in (Wu et al., 2010), indicating the proximity of two different proteins populations. PPI of 0 indicates no proximity (or no colocalization), and PPI of 1 indicates perfect proximity (or perfect colocalization); intermediate values give an estimate of the fraction of a given protein being in close proximity (or colocalize) with another one. Here the result of the proximity analysis is represented graphically with color-coded values and Venn diagrams as depicted on the top of the figure (details in Material and Methods). The analysis has been carried out on apico-basal (AB) or planar (PL) orientation images to minimize apparent colocalization due to overlapping in different planes; this is reported in the representative image of each experiment. (**A**) Proximity analysis for PATJ, PALS1, aPKC and PAR6β and corresponding representative images. Zoomed images (PATJ/aPKC and PALS1/PAR6β) illustrate the segregation of these proteins. (**B**) Proximity analysis for CRB3A and the other polarity proteins. (**C**) Proximity analysis for PAR3 with PALS1, aPKC and occludin. (**D**) Control experiment with PATJ labelled with an Alexa 532 secondary antibody and an Alexa 568 tertiary antibody. The number of junctions used in quantification and the details of the analysis are specified in Material and Methods. Scale bars: 1 µm.

First, we found that some of the proteins colocalize strongly: PALS1 and PATJ seem to reside in the same clusters, similarly to aPKC and PAR6β that also colocalize strongly, presumably in both cases forming a complex, as the literature suggests (Joberty et al., 2000; Lin et al., 2000; Roh, Makarova, et al., 2002) (Figure 3A). Surprisingly, we found PALS1-PATJ and aPKC-PAR6β well segregated from each other when we observed them in the planar orientation. They sometimes appeared as alternating bands along the junction with a spatial repeat in the range of 200 nm to 300 nm (zooms in Figure 3A). In some cases, these bands seemed formed by clusters facing each other in neighboring cells, indicating a potential coordination of polarity protein organization between adjacent cells. Second, we found that only a minority of CRB3A colocalized with any of the other polarity proteins (Figure 3B). These observations are also surprising, because CRB3A has been reported to strongly interact both with PALS1 and PAR6 (Hayase et al., 2013; Lemmers et al., 2004; Li et al., 2014; Makarova et al., 2003). This could mean that these interactions are mostly transient or that they are not prominent in the TJ area. This result questions the stability and functional cellular meaning of the canonical Crumbs-PALS1-PATJ complex and of the CRB3-PAR6 interaction. Finally, when localizing PAR3 along with PALS1 or aPKC, we found that PAR3 is hardly found with either of these proteins (Figure 3C). These data show that PAR3, aPKC and PAR6β do not associate in a static complex as it has been suggested in several non-mammalian models (Afonso & Henrique, 2006; Harris & Peifer, 2005; Morais-de-Sá et al., 2010; Rodriguez et al., 2017). It appears, in our conditions, that aPKC and PAR6β are likely linked in the apical TJ region, whereas PAR3 is mostly not associated to them. Again, it is possible that the interaction between PAR3 and PAR6β-aPKC is mostly transient or that it is not relevant in the TJ area. We conclude that PAR3 is mostly isolated from other polarity proteins at the TJ, and that PALS1-PATJ, PAR6β-aPKC and CRB3 form three spatially separated entities in the apical region of the TJ.

### PATJ localization in the TJ region with electron tomography

In the generally accepted description of the canonical Crumbs complex, PALS1 binds to the transmembrane protein CRB3A and PATJ binds to PALS1 (Roh, Makarova, et al., 2002). Therefore, PALS1 and PATJ are thought to be in close vicinity of the membrane since CRB3A is a short transmembrane protein. Moreover, it was proposed that PATJ links CRB3A-PALS1 to the TJ area (Michel et al., 2005) because of PATJ direct interaction with the TJ protein ZO-3 and Claudin1 (Roh, Liu, et al., 2002). Our protein-proximity analyses, however, raise the question of whether PALS1/PATJ interact with CRB3A in the TJ region (Figure 3), and our localization of PATJ with STED suggests that most PATJ proteins are often too far from the TJ to interact with this structure (Figure 1 and 2). Therefore, to obtain a more complete understanding of PATJ localization in the TJ region, we observed PATJ with electron tomography using immunogold labelling in Caco-2 cells (Figure 4).

**Figure 4.**
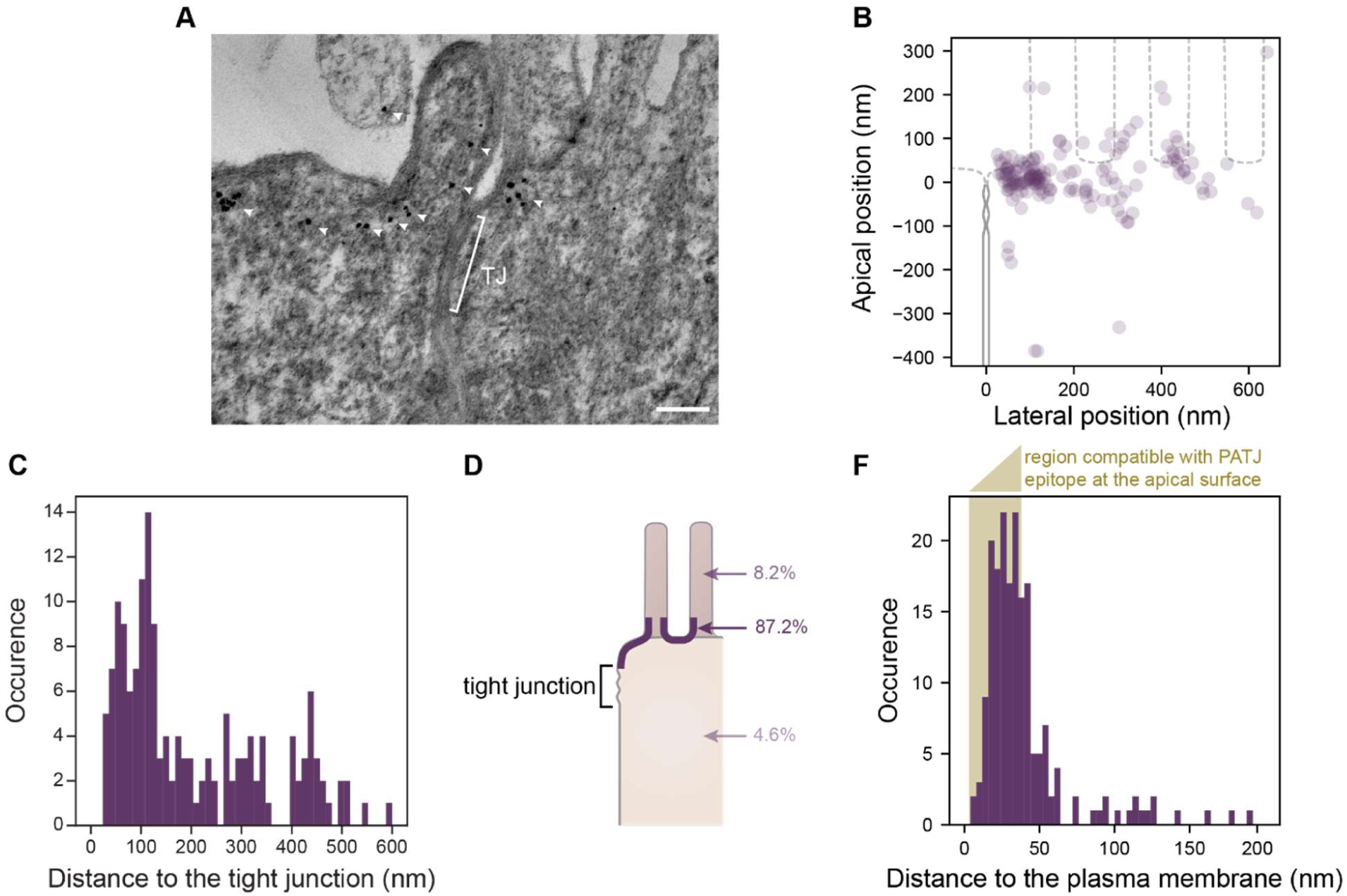
Electron tomography shows that PATJ localize as clusters at the plasma membrane apically of the TJ in Caco-2 cells. (**A**) Representative image of PATJ labelled with gold particles (arrowheads pointing at single particles or cluster of particles). Bracket with TJ indicate the tight junction. Minimum intensity projection of a 150 nm thick tomogram, scale bar: 100 nm. (**B**) Localization of gold particles labelling PATJ with respect to the TJ both in the apico-basal and lateral directions. (**C**) Distance between the center of gold particle labels and the TJ. (**D**) Summary of gold particles localization in the microvilli, in the vicinity of the plasma membrane and the cytoplasm. (**E**) Distance between gold particles and the apical surface. In amber, the region of distances compatible with PATJ epitope being at the apical surface, between 3 nm (radius of gold particles) and 37 nm (size of the primary and gold-labelled secondary antibody combination added with the presumed size of PALS1 (Li et al., 2014)).

Consistent with what we observed with STED, we often found PATJ organized in clusters apical of the TJ (Figure 4A). We started by quantifying PATJ position with respect to the TJ, using as a reference the most apical part of the TJ (defined morphologically as the most apical position of contact between neighboring cells plasma membranes) (Figure 4B). We found that most PATJ proteins were about 80 nm away from the TJ (Figure 4C). Although PATJ molecular structure is not known, given its sequence including multiple potent unstructured domains, it is likely to be a globular protein, which size cannot fill the 80 nm gap we find, with the nanometer-sized proteins of the TJ. Therefore, our data suggest that most PATJ molecules do not interact directly with TJ proteins. We found instead most PATJ proteins close to the apical membrane and that only a small fraction was present in microvilli or in the cytoplasm (Figure 4D). Previous observations that PATJ associate with ZO-3 or Claudin1 might depend on the cellular state or these interactions could be transient.

CRB3A is thought to anchor PALS1 and PATJ to the plasma membrane. However, given our results showing a minor colocalization of PATJ and PALS1 with CRB3A, it is unlikely to be the case for most PALS1 and PATJ molecules. Therefore, the localization of PATJ close to the apical membrane led us to wonder whether PATJ together with PALS1 could be associated with the apical plasma membrane via interactors that remain to be discovered. Thus, we measured the distance of the immunogold label of PATJ to the plasma membrane (Figure 4E) and found that the distance of the gold label is in most cases compatible with the association of PATJ and PALS1 with the apical plasma membrane (123/169 ≈ 73% of gold particles were less than 38 nm away from the plasma membrane, corresponding to the size of the primary and gold-labelled secondary antibody combination added with the size of PALS1). We conclude that PATJ and PALS1 are likely to be anchored to the apical membrane not by CRB3A but by yet unknown apical membrane proteins.

### Organization of PATJ-PALS1, PAR6β-aPKC and the actin cytoskeleton

Because polarity proteins play a key role in epithelial organization, we wondered how these proteins were organized with respect to the actin cytoskeleton. When labelling aPKC together with the actin-staining phalloidin, we found with 3D STED that aPKC labelled actin in microvilli localized in the direct vicinity of the junction (Figure 5A and supplement movie 1). Since we observed that PALS1-PATJ and PAR6β-aPKC complexes localize above the TJ in an alternated pattern (Figure 3A), and because of PATJ localization (Figure 4D), it appeared that the labelling pattern corresponded to the alternation of PAR6β-aPKC at microvilli and PALS1-PATJ at the plasma membrane in between microvilli. As a result, the patterns of PALS1-PATJ and PAR6β-aPKC complexes seem to follow the organization of actin just above the TJ.

**Figure 5.**
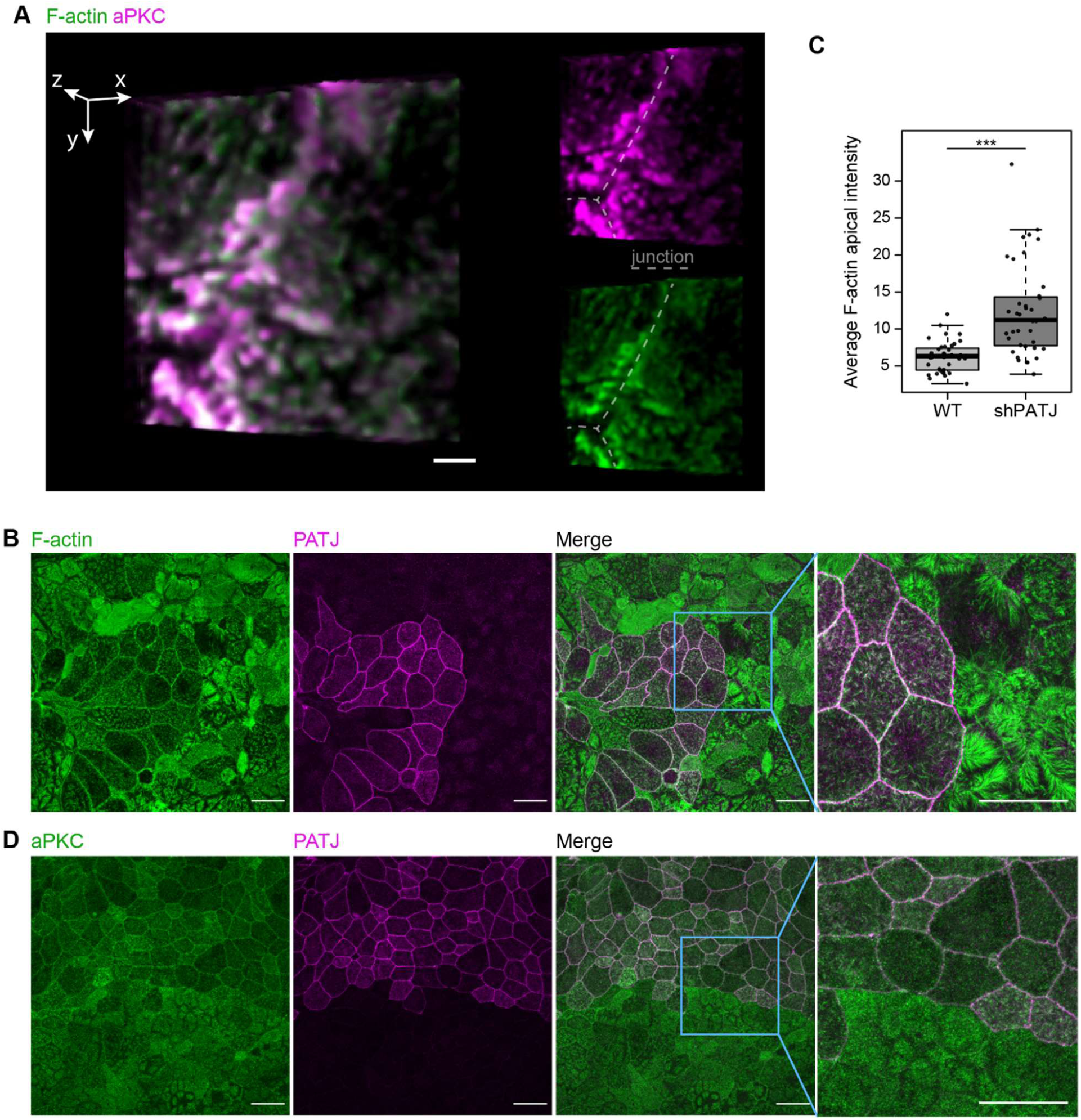
Actin organization and PATJ-PALS1 / aPKC-PAR6β complexes. (**A**) Localization of aPKC with respect to F-actin. 3D rendering of a 3D STED image stack extracted from the supplemental movie 1. The orientation of this image is slightly titled from the planar orientation to ease visualization. aPKC in magenta, phalloidin staining in green. Scale bar (for the merged color image) 1 µm. (B-D) Effect of PATJ depletion on the organization of apical actin and aPKC. To evaluate the effect of PATJ depletion, we used a mix of WT and KD PATJ Caco-2 cells. (**B**) Effect of PATJ depletion on the actin organization at the apical surface (PATJ in magenta, phalloidin staining in green). (**C**) Quantification of the average apical intensity in WT versus shPATJ cells. Junctions were excluded from the analysis. Because of the non-normality of data, we used the Wilcoxon rank sum test to test for the difference of median between WT and shPATJ samples average apical phalloidin staining intensity; we obtained p-value = 7.9e-09. (**D**) Effect of PATJ depletion on aPKC organization at the apical surface (PATJ in magenta, aPKC staining in green). Scale bars 20 µm.

How PALS1-PATJ and PAR6β-aPKC complexes interact with actin is unknown. As an attempt to uncover a potential role of these complexes in the organization of actin in the area, we used a downregulated PATJ stable clonal line of Caco-2 cells (clone 4 from (Michel et al., 2005)) and mixed them with WT Caco-2 cells to compare protein localizations in both cell types grown in the same conditions. We and others have already shown that the depletion of PATJ impairs the TJ and depletes both PALS1 and CRB3 from the TJ region (Michel et al., 2005; Shin et al., 2005). In the apical part of WT Caco-2 cells, actin is present in microvilli and in an apparent belt at the adherens junction level (Mangeol et al., 2019). In contrast, we found in PATJ KD cells that the distribution of apical actin was strongly affected (Figure 5B). In downregulated PATJ cells, the intensity of apical actin was doubled on average in comparison to WT cells (Figure 5C). Moreover, while the actin belt was easy to identify in WT cells, it was sometimes difficult to discern it in PATJ downregulated cells. Similarly, aPKC intensity was increased towards the apical membrane in many PATJ knock-down cells (Figure 5D). These results show that PATJ influences the regulation of the actin cytoskeleton organization in the apical region of Caco-2 cells.

## Discussion

In this study, we have systematically localized polarity proteins with super-resolution microscopy in epithelial cells. We observed endogenous PAR3, aPKC, PAR6β, PATJ, PALS1 and CRB3A in human intestine and Caco-2 cells, and PAR3 and PALS1 in mouse intestine. We found the following (Figure 6). (**1**) All these polarity proteins organize as submicrometric clusters concentrated in the TJ region. PAR3 localizes at the TJ, aPKC and PAR6β localize at the tight junction level, but mostly apically of the TJ, while PATJ, PALS1 and CRB3A are apical of the TJ (Figures 1,2). (**2**) PAR6β-aPKC and PATJ-PALS1 form two pairs that are often respectively found in the same clusters (Figure 3A), strongly indicating that these respective proteins form a stable and major complex in this region of the cells (*i.e* the PAR6-aPKC complex and the PALS1-PATJ complex). (**3**) Unexpectedly, PALS1-PATJ and PAR6β-aPKC clusters are segregated from each other (Figure 3A). Our data suggest that the PAR6β-aPKC complex is localized at the base of the first row of microvilli in the direct vicinity of the TJ, whereas PALS1-PATJ is localized between the TJ and these microvilli, as well as in between these microvilli (Figure 4,5). This direct link between actin organization and polarity protein localization led us to probe the effect of PATJ on actin organization. (**4**) We found that PATJ regulates the organization of filamentous actin in the area, as the depletion of the PATJ affects both microvilli and the apical actin belt (Figure 5). (**5**) CRB3 shows little association with any of the other polarity proteins (Figure 3B), questioning how PALS1-PATJ and PAR6β-aPKC are mechanistically recruited to the plasma membrane and localized to the apical surface.

**Figure 6.**
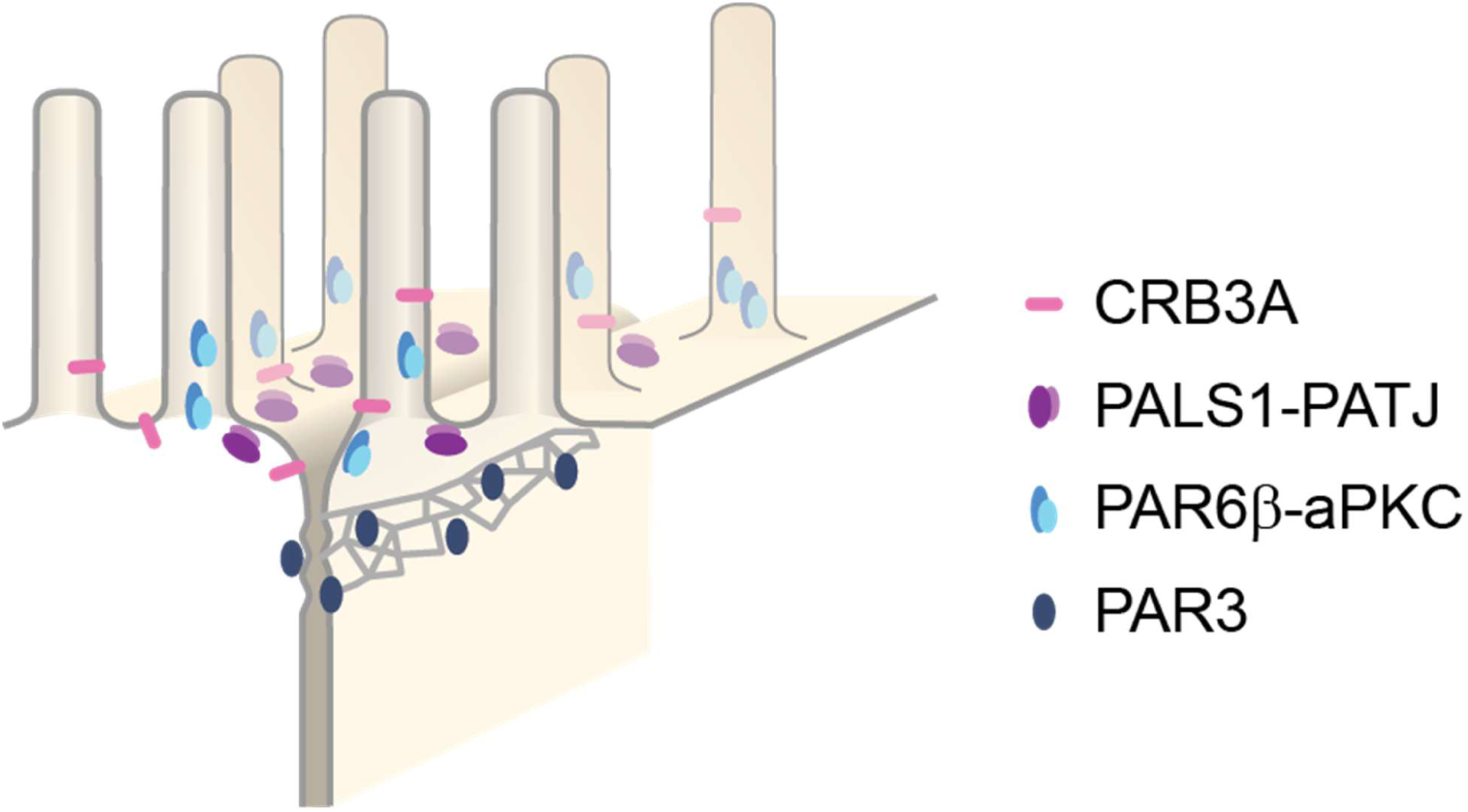
Organizational model of polarity proteins in the TJ region.

Previous studies were largely based on biochemical approaches. The first interactions found defined canonical polarity protein complexes, while subsequent studies highlighted the numerous potential interactions that can be found with such approach, between polarity proteins of different complexes, (Assémat et al., 2008; Bhat et al., 1999; Hurd et al., 2003; Joberty et al., 2000; Lemmers et al., 2004; Lin et al., 2000; Makarova et al., 2003; Roh, Makarova, et al., 2002) as well as between polarity proteins and other interactors (Chen & Macara, 2005; Itoh et al., 2001; Médina et al., 2002; Michel et al., 2005; Roh, Liu, et al., 2002; Takekuni et al., 2003; Tan et al., 2020). Altogether these studies provide a complex potential model of molecular interactions. However, in most cases, we do not know to what extent and where these interactions do occur in cells and whether they are transient or permanent. Notably, most of these previous studies used overexpression to identify the interactors of a given protein; this methodological limitation might have introduced false-positive in some cases. In an attempt to reduce the complexity of the current view, our study proposes a snapshot in the mature intestinal epithelia to simultaneously localize endogenous proteins two-by-two with unprecedented spatial resolution. Our results may bring a new light to the understanding of polarity proteins interactions, as it defines polarity complexes as they occur in the apical epithelial junction region. In particular, we question the existence of the canonical Crumbs and PAR complexes as previously described and propose that only PAR6β-aPKC and PALS1-PATJ can be defined as major structural complexes. The other numerous possible interactions that have been claimed previously may exist transiently and our approach cannot rule out that they occur at other locations in the cell, but it questions their relevance to the understanding of the epithelia cell junction.

The interaction between PAR3, PAR6 and aPKC is key to epithelial polarization (Horikoshi et al., 2009; Joberty et al., 2000) but the permanence of these interactions has been discussed in the past. In mammalian epithelial cells, PAR3, PAR6 and aPKC have been thought to interact at apical junctions as these proteins concentrate there, but only PAR6 and aPKC are found at the apical surface (Martin-Belmonte et al., 2007; Satohisa et al., 2005). Moreover in a few non-mammalian systems, PAR3 was observed as segregated from PAR6 and aPKC at epithelial apical junctions: when observed with confocal microscopy, PAR3 is clearly basal of PAR6 and aPKC in the apical junctions of *Drosophila melanogaste*r embryos during cellularization (Harris & Peifer, 2005), as well as in chick neuroepithelial cells (Afonso & Henrique, 2006). Our data suggest that the segregation of PAR3 from PAR6-aPKC is likely to be a conserved principle of organization in polarized epithelia. Even if the interaction of PAR3 with PAR6-aPKC is central to polarization, it is not permanent. The mechanistic basis for the transient character of the interaction between PAR3 and PAR6-aPKC in mammalian epithelia may be similar to the Cdc-42-dependant mechanisms found in *Drosophila melanogaster* (Morais-de-Sá et al., 2010) or *Caenorhabditis elegans* (Rodriguez et al., 2017).

Our finding that PAR3 localizes at the TJ confirms previous observations using electron microscopy in rat small intestine (Izumi et al., 1998) and MDCK cells. One recent study found a small fraction of PAR3 at the level of the adherens junction (Tan et al., 2020). Even though STED allows for much larger volume to be probed compared to electron microscopy, we did not observe PAR3 basal of the TJ. The localization of PAR3 may depend on the cell type as well as its maturation state, but interestingly PAR3 is never found in the region apical of the TJ, where we find the other polarity proteins.

Because CRB3 is a transmembrane protein and that several studies reported its interaction with PALS1, it was thought to anchor PALS1 and PATJ to the apical membrane (Makarova et al., 2003; Roh, Makarova, et al., 2002). Similarly, it is suggested in *Drosophila melanogaster* that Crb recruits PAR6 and aPKC to the apical membrane (Morais-de-Sá et al., 2010). Our study suggests that the recruitment of PALS1, PATJ, PAR6β, and aPKC to the plasma membrane is unlikely to be due to CRB3A, because CRB3A poorly colocalizes with these proteins. Nevertheless, our data suggest that PALS1-PATJ are localized at the plasma membrane, perhaps confined in this area by another set of interactors to be uncovered. This last observation is likely to be similar for PAR6-aPKC. We cannot rule out both for PALS1-PATJ and PAR6β-aPKC that the interaction with CRB3A could be transient, and that this transient interaction would be sufficient to localize these proteins complexes in the apical surface area.

The importance of polarity proteins for the epithelial organization point at the fact that these proteins are likely to play a key role in the organization of the cytoskeleton. Several proteins having a role in actin regulation have been shown to interact with polarity proteins (Bazellières et al., 2018; Médina et al., 2002), but how polarity protein could influence actin organization is largely unknown. The correlation of organization between the actin cytoskeleton and PAR6-aPKC and PALS1-PATJ clusters points at a potentially structural role of these proteins to the cytoskeleton organization. These findings call for further investigations, including functional and structural approaches.

In this study, we define endogenous polarity protein organization and how polarity protein are likely to interact. The early concept of polarity protein complexes introduced by biochemical studies is impractical today because of the very large number of potential interactions between proteins discovered. Additionally, it omits important features, such as the dynamics of interaction as well as their reality in relation to cell sub-regions. Our study proposes a snapshot of the polarity organization in mature intestinal epithelial cells that calls for novel, more dynamic definition of interactions between polarity proteins and associated proteins that will be needed to uncover the mechanistic basis of cell apico-basal polarization.

## Materials and Methods

### Cell culture

A clone of Caco-2 cells, TC7, was used in this study because differentiated TC7 cells form a regular epithelial monolayer (Chantret et al., 1994). Cells were seeded at a low concentration of 10^5^ cells on a 24 mm polyester filter with 0.4 µm pores (3450, Corning inc., Corning, NY). Cells were maintained in Dulbecco’s modified Eagle’s minimum essential medium supplemented with 20% heat-inactivated fetal bovine serum and 1% non-essential amino acids (Gibco, Waltham, MA), and cultured in 10% CO_2_/90% air. The medium was changed every 48 hours.

### Sample preparation for immunostaining

#### Human sample preparation

Human intestine biopsies were obtained under the agreement IPC-CNRS-AMU 154736/MB. Intestinal samples were fixed in paraformaldehyde (PFA 32%, Fischer Scientific) 4% in phosphate buffer saline (PBS, Gibco, Waltham, MA) for 1 hour at 20°C. Biopsies were embedded in optimal cutting temperature compound (OCT compound, VWR) and frozen in liquid nitrogen.

#### Mouse sample preparation

Mouse intestine samples were obtained following ethical guidelines. After washing with PBS intestinal samples were fixed in PFA 4% in PBS for 20 minutes at room temperature. Samples were then embedded in OCT compound and frozen in liquid nitrogen.

#### Cell culture preparation for optical microscopy

Cells were washed in PBS and then fixed in PFA 4% in PBS for 20 minutes at room temperature. When apico-basal orientation observations were needed, cells were sectioned along the apico-basal axis. Prior sectioning, cells were embedded in OCT compound and frozen in liquid nitrogen.

#### Samples sectioning

When needed, samples were sectioned with a cryostat (Leica CM 3050 S, Leica Biosystems, Germany). 10 µm sections were transferred to high precision 1.5H coverslips (Marienfeld, Germany) previously incubated with Poly-L-lysine solution (P-4832, Sigma-Aldrich, St. Louis, MO).

### Immunostaining for optical microscopy

Intestinal sections and cultured cells were prepared similarly. Intestinal sections were permeabilized in 1% SDS (Sigma-Aldrich) in PBS for 10 minutes. In cultured cells, 10 minutes permeabilization was achieved with 1% SDS in PBS for CRB3A antibody, as well as PAR6β and aPKC antibodies when used in combination with tight junction markers; otherwise all other protein labelling were using 1% Triton X100 (Sigma-Aldrich) in PBS permeabilization for 10 minutes. After washing with PBS, samples were saturated with 10% fetal bovine sera (Gibco) in PBS (“saturation buffer”) over an hour at room temperature. Primary antibodies were diluted in the saturation buffer and incubated overnight at 4°C. In more details: rabbit anti-ZO-1 (1/500, 61-7300, Invitrogen), mouse anti-occludin (1/500, 331500, Invitrogen), mouse anti-E-cadherin (1/500, 610181, BD Biosciences), rabbit anti-PAR3 (1/200, 07-330, Sigma-Aldrich), rabbit anti-PAR6β (1/200, sc-67393, Santa-Cruz), rabbit anti-PKCζ (1/200, sc-216, SantaCruz), mouse anti-PKCζ (1/200, sc-17781, SantaCruz), chicken anti-PALS1 (1/200, gift of Jan Wijnholds (Kantardzhieva et al., 2005)), rabbit anti-PATJ (1/200, (Massey-Harroche et al., 2007; Michel et al., 2005)), rat anti-CRB3A (1/50 MABT1366, Merck). Secondary antibodies were incubated 1 hour at room temperature. Alexa Fluor 568 conjugated to antibodies raised against mouse, rabbit and rat and Alexa Fluor 532 conjugated to antibodies raised against mouse and rabbit (Invitrogen) were used at 1/200 dilution in the saturation media. Phalloidin Alexa Fluor A532 (Invitrogen) was mixed with secondary antibodies and used at 1/100 dilution. After each incubation, samples were rinsed 4 times with PBS. Samples were finally mounted in Prolong Gold antifade mountant (Invitrogen) at 37°C for 45 minutes.

### STED microscopy

Images of samples were acquired with a STED microscope (Leica TCS SP8 STED, Leica Microsystems GmbH, Wetzlar, Germany), using a 100X oil immersion objective (STED WHITE, HC PL APO 100x/1.40, same supplier). Two-color STED was performed with Alexa Fluor 532 excited at 522 nm (fluorescence detection in the 532-555 nm window), and Alexa Fluor 568 excited at 585 nm (fluorescence detection in the 595-646 nm window). To minimize the effect of drifts on imaging, both dyes were imaged sequentially on each line of an image and depleted using the same 660 nm laser. Detection was gated to improve STED signal specificity.

### Cultured cell preparation for electron microscopy

Cells were washed in PBS and then fixed in PFA 4% in PBS for 20 minutes at room temperature. After rinsing with PBS, cells were put into a sucrose gradient to reach 30% sucrose overnight. Cells were then frozen in liquid nitrogen and immediately thawed at room temperature. Immunostaining was carried out without permeabilization step, directly with primary antibodies (rabbit anti-PATJ 1/100, for 3 hours at room temperature). After washing steps, cells were incubated with secondary antibody carrying 6 nm gold particles (goat anti-rabbit 1/20, 806.011, Aurion, The Netherlands). A tertiary antibody was used to observe where gold particles were localized on a macroscopic level (Alexa 568 conjugated donkey anti-goat 1/200 from Invitrogen, for 1 hour at room temperature).

Cells were then prepared specifically for electron microscopy. They were fixed in 2.5% glutaraldehyde, 2% PFA, 0.1% tannic acid in sodium cacodylate 0.1M solution for 30 minutes at room temperature. After washing steps, cells were post-fixed in 1% osmium in sodium cacodylate 0.1M solution for 30 minutes at room temperature and contrasted in 2% uranyl acetate in water solution for 30 minutes at room temperature. Cells were then dehydrated in ethanol and embedded in Epon epoxy resin.

### Electron microscopy

Cells were observed with a transmission electron microscope, FEI Tecnai G2 200 kV (FEI, The Netherlands), in an electron tomography mode. Tomograms were reconstructed using the Etomo tool of the IMOD software.

### Data analysis

#### Analysis of protein density

To quantify the density and positions of polarity proteins with respect to tight junction markers, we used custom-made ImageJ macros and Python programs. In each case the reference protein was a tight junction protein (ZO-1 or occludin) that was localized precisely, defining a reference position along the junction from which intensity measurement was done. For planar orientations, the reference was the maximum intensity of the tight junction marker along of the junction; intensity measurements consisted in getting the intensity profiles of proteins perpendicular to the junction, all along the junction. For apico-basal orientations, we measured intensity profiles on the apico-basal axis, all along the junction. On a given profile, the reference was taken at the most apical point where the tight junction marker intensity was a third of its maximum intensity; the reason for this choice is that tight junctions spread along the apico-basal axis tended to vary up to three-fold from one cell to another and this definition of the reference allowed us to define a reproducible apical edge of the tight junction. In the process, we used bilinear interpolation to obtain sub-pixel quantification. Results of analyses were then normalized for intensity for each junction to avoid junction-to-junction intensity variation. Because we used a reference protein for each junction, we could then align all results based on the reference position of the reference protein and pool all results into a single protein density plot.

### Protein-proximity analysis

The principle of quantification of protein-proximity was proposed in (Wu et al., 2010). The authors of this method observed that the autocorrelation of a given image or the cross-correlation between two images coming from two different channels showed a peak at its center. The ratio of amplitude between the peaks of the cross-correlated and autocorrelated images gave a good estimate of protein proximity, which they coined the protein-protein proximity index. This index is similar to more classical colocalization coefficients, but we found that the method of (Wu et al., 2010) was well suited for proteins distributed along a junction.

In practice, we extracted junctions from two-color images, restricting the analysis to a band of 400 nm centered on the reference given by the tight junction (as defined in the previous paragraph). As we found the analysis to be dependent on orientation, when planar orientation was used, we excluded junctions that were not straight. All extracted junctions of a given protein pair to be examined were then concatenated into one large two-channel image on which we achieved autocorrelation and cross-correlation analysis (autocorrelation is achieved on each channel, and cross-correlation is achieved with both channels). We extracted the amplitude of peaks obtained in each of the autocorrelated and cross-correlated images as proposed in (Wu et al., 2010). Therefore, when analyzing protein 1 and protein 2 proximity, we obtain the amplitude A_1_ and A_2_ from the autocorrelation of images of protein 1 and protein 2 respectively, and the amplitude C_12_ from the cross-correlation analysis. One evaluates the fraction of protein 1 colocalizing with protein 2 with the protein-protein proximity index P_1_ = C_12_/A_2_, and the fraction of protein 2 colocalizing with protein 1 with the protein-protein proximity index P_2_ = C_12_/A_1_.

In figure 2 we color coded the values of these indices. In order to obtain an absolute representation of these values, we additionally used Venn diagrams to represent graphically for each protein the fraction of colocalizing and non-colocalizing protein.

### Number of junctions or cells used in the analysis

**Figure 1 and 2**

Number of junctions used in the analysis. Pl: planar, AB: apico-basal

| Label | PAR3 Occl | | aPKC Occl | | PAR6β Occl | | PATJ Occl | | PALS1 ZO-1 | | CRB3A ZO-1 | |
| --- | --- | --- | --- | --- | --- | --- | --- | --- | --- | --- | --- | --- |
| Sample | PI | AB | PI | AB | PI | AB | PI | AB | PI | AB | PI | AB |
| Caco-2 | 29 | 8 | 16 | 9 | 18 | 11 | 33 | 8 | 12 | 9 | 16 | 12 |
| Human | 25 | 7 | 16 | 10 | 12 | 12 | 47 | 16 | 23 | 10 | 12 | 8 |
|  | PAR3 Occl |  |  |  |  |  |  |  | PALS1 Occl |  |  |  |
| Mouse | 10 | 9 |  |  |  |  |  |  | 10 | 8 |  |  |

**Figure 3**

| Label | PATJ<br>PALS1 | PATJ<br>aPKC | PALS1<br>aPKC | PALS1<br>PAR6β | PAR6β<br>aPKC | CRB3A<br>PALS1 | CRB3A<br>PATJ |
| --- | --- | --- | --- | --- | --- | --- | --- |
| Number of junctions | 21 | 42 | 36 | 27 | 31 | 25 | 25 |
| Label | CRB3A aPKC | CRB3A<br>PAR6β | CRB3A<br>PAR3 | CRB3A<br>ZO-1 | PAR3<br>PALS1 | PAR3<br>aPKC | PAR3<br>OCLN |
| Number of junctions | 17 | 18 | 9 | 16 | 9 | 27 | 15 |
| Label | PATJ-Alexa568<br>PATJ-Alexa532 |  |  |  |  |  |  |
| Number of junctions | 15 |  |  |  |  |  |  |

**Figure 4**

Tomograms of 300 nm in thickness of 12 junctions were used to extract the position of 169 gold particles labelling PATJ proteins.

**Figure 5**

Number of cells used in Figure 4B quantification: 39 WT cells and 40 PATJ downregulated cells (from one sample, in two different areas).

## Acknowledgments

We would like to thank Flora Poizat for human biopsies and Le Bivic and Lenne groups for discussion. Biopsies were obtained with the agreement IPC-CNRS-AMU 154736/MB. Funding: We acknowledge the IBDM imaging facility, member of the national infrastructure France-BioImaging supported by the French National Research Agency (ANR-10-INBS-04). PM was supported by ITMO Cancer (Plan Cancer), Ligue nationale contre le Cancer and the French National Research Agency (ANR-T-JUST, ANR-17-CE14-0032). The project developed in the context of the LabEx INFORM (ANR-11-LABX-0054) and of the A*MIDEX project (ANR-11-IDEX-0001-02), funded by the “Investissements d’Avenir” French Government program.

## Author contributions

P.M. performed the experiments and analyzed the data. D.M-H. assisted in performing most experiments. F.R. prepared samples for electron microscopy and acquired electron tomograms. P.M., A.L.B and P-F.L. designed the experiments and acquired financial support for the project. All authors discussed the results and contributed to the manuscript.

